# Convergence of clinically relevant manipulations on dopamine-regulated prefrontal activity underlying stress-coping responses

**DOI:** 10.1101/2021.03.20.436282

**Authors:** Scott A. Wilke, Karen Lavi, Sujin Byeon, Vikaas S. Sohal

## Abstract

**Background:** Depression is a pleiotropic condition that can be produced or ameliorated by diverse genetic, environmental, and pharmacological manipulations. In this context, identifying patterns of circuit activity on which many of these manipulations converge would be important, because studying these patterns could reveal underlying biological processes related to depression and/or new therapies. In particular, the prefrontal cortex and dopaminergic signaling have both been implicated in depression. Nevertheless, how dopamine influences disease-relevant patterns of prefrontal circuit activity remains unknown.

**Methods:** We used calcium imaging in brain slices to identify depression-relevant patterns of activity in prefrontal microcircuits, and measure how these are modulated by dopamine D2 receptors (D2Rs). Then, we used optogenetic and genetic manipulations to test how dopamine and D2Rs contribute to stress-coping behavior in a paradigm commonly used to assay how manipulations promote or ameliorate depression-like states.

**Results:** Patterns of correlated activity in prefrontal microcircuits are enhanced by D2R stimulation as well as by two mechanistically distinct antidepressants: ketamine and fluoxetine. Conversely, this D2R-driven effect was disrupted in two etiologically distinct models of depression: a genetic susceptibility model and chronic social defeat. Phasic stimulation of dopamine afferents to prefrontal cortex increased effortful responses to tail suspension stress. Conversely, deleting prefrontal D2R receptors reduced the duration of individual struggling episodes.

**Conclusions:** Correlated prefrontal microcircuit activity represents a point of convergence for multiple depression-related manipulations. Prefrontal D2Rs enhance this activity. Through this mechanism, prefrontal dopamine signaling may promote network states associated with antidepressant actions that manifest as effortful responses to stress.

## INTRODUCTION

Numerous mechanistic studies have implicated prefrontal circuits in human depression or depression-like states in animal models (1–6). Indeed, directly modulating activity in prefrontal circuits can relieve many symptoms associated with clinical depression (7–9). A wide range of genetic and environmental factors influence depression susceptibility (10), and a similarly broad range of interventions, including medications, electrical stimulation, and exercise, can alleviate depressive symptoms. Here, we take advantage of this heterogeneity by searching for convergent microcircuit processes engaged by these disparate factors. Seeking convergence is a tactical approach for filtering pleiotropic changes in neural circuits in order to identify those mechanisms which are most likely to have broad clinical relevance. We adopted a multi-step process. First, we used a relatively simple slice calcium imaging assay (Fig. 1) to rapidly screen for convergent changes in circuit activity elicited by several factors with clinical relevance to depression. These include mechanistically distinct antidepressant medications, as well as mice which model genetic and environmental factors related to depression-like states and depression susceptibility or resilience. We identified a signature of prefrontal circuit activity that is consistently modulated in opposing directions by manipulations which either exacerbate or ameliorate aspects of depression or depression-like phenotypes. Furthermore, we find that activating dopamine D2 receptors (D2Rs) enhances pattern of activity which are normally recruited by antidepressant treatments. This ability of D2Rs to recruit antidepressant-related patterns of activity is disrupted in two etiologically distinct models of depression-like states. Based on this, we predict that driving or disrupting signaling through D2Rs should promote or suppress behaviors that are associated with antidepressant response, respectively. We then tested this prediction using the tail suspension test (TST) to measure effects of optogenetically evoking prefrontal dopamine release or knocking out prefrontal D2Rs.

**Fig. 1.**
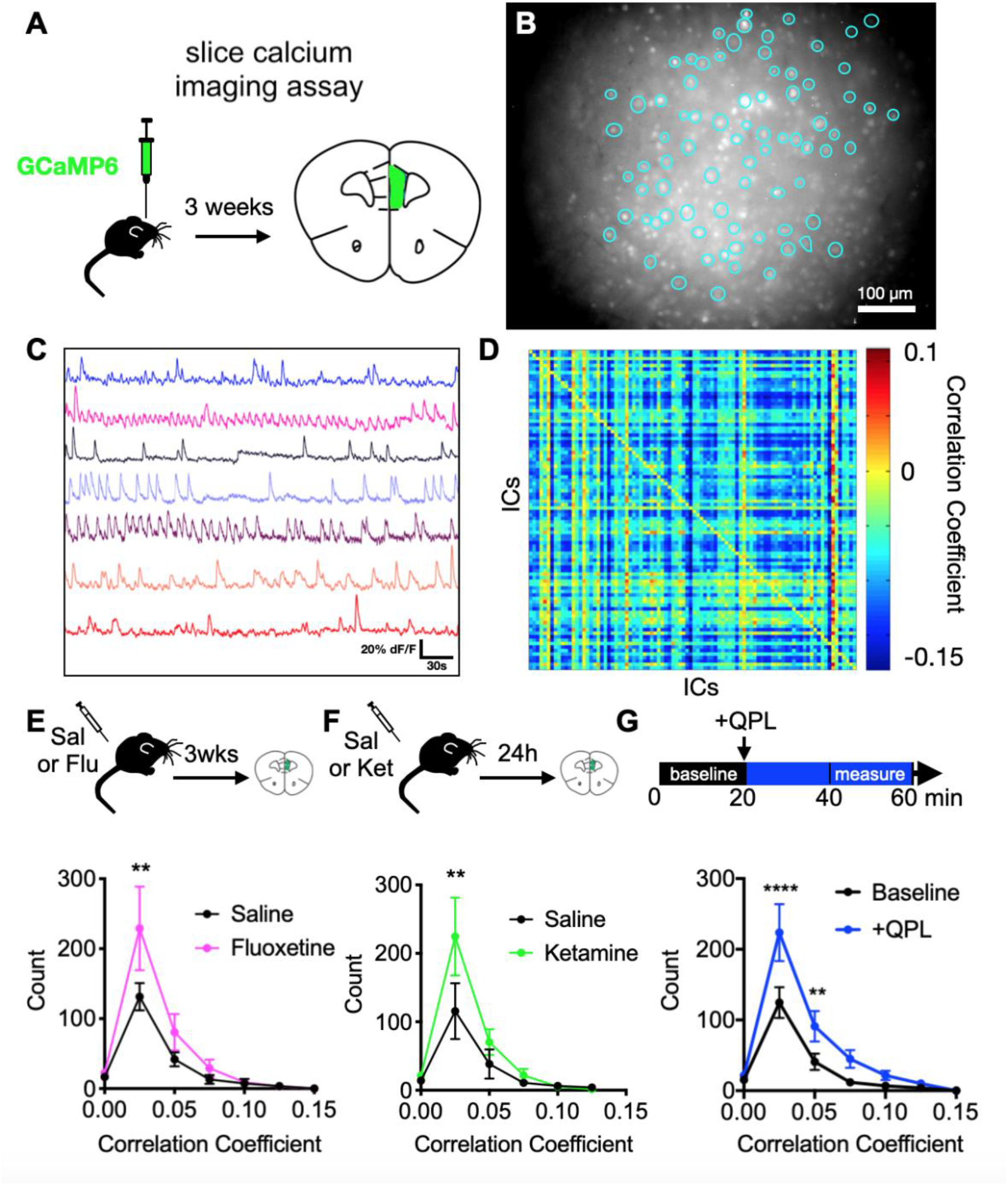
Antidepressants and the dopamine D2 agonist quinpirole increase positively correlated activity in mPFC acute slices. (**A**) Schematic of GCaMP6f injection into mPFC. (**B**) Example field of view for Ca^2+^ imaging with regions of interest (ROI) outlined. (**C**) GCaMP signals from 8 example cells. (**D**) Heatmap showing correlations between 85 cells in baseline conditions. IC, independent component. (**E–G**) Experimental design (top) and quantification of effects on the distribution of positive correlations in mPFC network activity (bottom) for (**E**) Fluoxetine [*F*_Interaction_(6,133)=1.64, *P*=0.14, *F*_Drug_(1,133)=4.68, *P*=0.032], (saline, n=10; fluoxetine, n=11) (**F**) Ketamine [*F*_Interaction_(5,90)=1.88, *P*=0.11, *F*_Drug_(1,90s)=4.19, *P*=0.044], (saline, n=9; ketamine, n=8) and (**G**) Quinpirole [*F*_Interaction_(6,252)=2.92, *P*=0.009, *F*_Drug_(1,252)=16.52, *P*<0.0001], (n=19), 2-way ANOVA with Sidak post-hoc test. For this and all subsequent figures, statistics represent mean±SEM, \**P*<0.05, \*\**P*<0.01, \*\*\**P*<0.001, \*\*\*\**P*<0.0001.

## MATERIALS AND METHODS

All experiments were conducted in accordance with procedures established by the administrative panels on laboratory animal care at the University of California, San Francisco.

### Mouse lines, viral vectors and viral targeting/expression

For all experiments using wildtype mice, C57BL/6J derived lines were bred in house or ordered from Jackson Laboratories or Charles River. Dominant negative Disc1 mutant mice were generated by crossing B6-CamKII::TtA (JAX: 00310) mice with tetO-DISC1dn (JAX: 008790) to yield mice expressing dominant negative DISC1 in neocortical pyramidal cells (11) For slice experiments, wildtype or Disc1 mutant mice at P26-P35 were injected with 4x 150 nl of AAV5-CaMKII-GCaMP6f (UPenn Virus Core) at 4 depths (dorso-ventral (DV): −2.0, −2.25, −2.50, −2.75) at the following (millimeters relative to bregma) coordinates for mPFC: 1.7 anterior-posterior (AP) and 0.3 mediolateral (ML). We allowed between 2-3 weeks for viral expression before sacrificing mice and cutting live prefrontal brain slices for calcium imaging. For optogenetics targeting VTA-mPFC projections, mice were TH::Cre (line FI12 www.gensat.org) and a mixture of male and female mice were used. Cre-dependent expression was driven using previously described adeno associated virus (AAV5) containing DIO-ChR2-eYFP or DIO-eYFP expressed under the synapsin promoter (12). We stereotaxically injected 1.0 µl of 4-10 × 10^12^ vg/ml virus into right VTA of P56-70 TH::Cre mice using methods described previously (12,13). Coordinates relative to bregma in millimeters were −2.75 AP, ±0.5 ML, and −4.5 DV for all experiments. At least 8 weeks were allowed for expression and trafficking to VTA terminals in mPFC. For Drd2 deletion experiments, mice were Drd2loxP/loxP homozygous (JAX: 020631) (14) Localized deletion of Drd2 in mPFC was accomplished using AAV5 containing Cre-mCherry or mCherry expressed under synapsin promoter (UPenn Virus Core). Coordinates relative to bregma were +1.7 AP, ±0.3 ML, and −2.5 DV and we injected 750 nl of 3-7 × 10^12^ vg/ml virus bilaterally in P56-P70 wildtype mice of either sex. At least 4 weeks were allowed for expression/deletion before proceeding with behavior experiments.

For optogenetics experiments, mice were implanted with a 200 µm diameter, 0.22NA fiber optic cannulae (Doric Lenses) over right mPFC (1.7 AP, 0.35 ML, −2.25 DV). Implants were affixed onto the skull using Metabond dental cement (Parkell). Targeting of viral constructs/cannulas, and expression of channelrhodopsin in prefrontal terminals were confirmed via histology without immunostaining.

### Slice calcium imaging experiments

In all cases, 350-micron thick live coronal slices were prepared from mice between 2-4 weeks after injection. Slice preparation followed our previously described protocol (15), briefly, slices were cut and incubated for 10 minutes in N-methyl-D-glucamine (NMDG)-based recovery solution before being transferred to ACSF for the remainder of the recovery period. The NMDG-based solution was maintained at 32C and used to maintain the overall health of adult slices to ensure sufficient activity for analysis. Prior to imaging, slices were moved from the recovery solution to one with 2 µM carbachol to promote adequate activity in prefrontal microcircuits during imaging. For details see (15) GCaMP6f imaging was performed on an Olympus BX51 upright microscope with a 20x 1.0NA water immersion lens, 0.5x reducer (Olympus, Tokyo, Japan), and ORCA-ER CCD Camera (Hamamatsu Photonics, Hamamatsu, Japan). Illumination was delivered using a Lambda DG4 arc lamp (Sutter Instruments, Novato, California). Light was delivered through a 472/30 excitation filter, 495 nm single-band dichroic, and 496 nm long pass emission filter (Semrock, Rochester, New York). All recordings were at 32.5 +/-1C. All movies that were analyzed consisted of 36,000 frames acquired at 10 Hz (1 hour) with 4 × 4 sensor binning yielding a final resolution of 256 × 312 pixels. For quinpirole stimulation experiments, baseline activity was imaged for 20 minutes, at which point the solution was switched to one containing 20 µM quinpirole to stimulate dopamine D2 receptors. Light power during imaging was 100 to 500 µW/mm^2^. The Micro Manager software suite (v1.4, National Institutes of Health, Bethesda, Maryland) was used to control all camera parameters and acquire movies. Subtle drift in x or y dimensions was corrected using an image alignment plugin for ImageJ software package, called cvMatch_Template, which effectively eliminated drift(16). Any movies with significant drift that could not be corrected or that lacked significant amounts of activity were excluded from further analysis. Cortical layers were clearly delineated by GCaMP expression and the bulk of infected neurons were localized to layer 5, where we focused our imaging window.

### Signal Extraction and correlations

All analyses and signal extraction was performed using MATLAB (Mathworks) and methods used for performing signal extraction are identical to our previous slice calcium imaging work. We used a PCA/ICA approach modified from the published CellSort 1.1 toolbox (17) as previously described ((15,18), detecting signals from between 80-100 ROIs per slice (presumed neurons). In brief, the baseline fluorescence function, F0, was calculated for every trace using the mode of the kernel density estimate over a 100s rolling window, implemented via the MATLAB function ksdensity following the procedure outlined in ((19). All signal traces shown represent normalized versions of the (F-F0)/F0 trace. Raw fluorescence correlations were calculated based on these traces, as described in the text and previously (20). In brief, we sought to summarize the patterns of local microcircuit activity using the pattern of correlations between activity in different neurons. Inter-neuronal correlations were computed as the normalized vector dot product between a time series of GCaMP signals and a time series of derivatives (corresponding to a different neuron). In addition to correlation, we also calculated a p-value for how well activity in one ROI predicts fluctuations in the other. Comparisons were made between conditions for positive correlations with a p-value less than 0.01. We used shuffled datasets (in which each trace has been shifted over by a random amount) or scrambled datasets (in which new traces were formed from random linear combinations of the original traces) to show that this algorithm detects statistically-meaningful correlations, i.e., correlations not present in shuffled/scrambled datasets (20).

### Drug treatments

For slice experiments combined with an in vivo pharmacologic manipulation (fluoxetine or ketamine), drugs were diluted in 0.9% sterile saline and injected intraperitoneally. Mice treated with fluoxetine were administered daily doses at 20 mg/kg every day for 4 weeks. For ketamine experiments, we first tested a range of ketamine doses (3, 10 and 50 mg/kg) alongside saline controls injected intraperitoneally (7 week old wildtype mice). Mice were tested 24 hours later using the TST in order to assess antidepressant-like, dose-response relationships. Based on this validation dose-response curve, 10 mg/kg was selected as the dosage of ketamine used for subsequent slice experiments as this had the maximal effect on reduction of immobility in the TST. For ketamine slice experiments, littermate pairs were injected together – one with 10 mg/kg ketamine and one with saline as a control – and slices were cut and imaged from both mice the next day (∼24 hours post-injection).

### Behavior experiments

Mice were housed in reversed 12-h light/dark cycles, and all experiments were performed during the light portion of the cycle. After sufficient time for surgical recovery and viral expression, mice underwent multiple rounds of habituation. The testing room was illuminated at 150 lux, and mice were first habituated to the behavioral testing area for 30 minutes prior to the beginning of any further handling each day. Mice were then habituated to touch with at least 3 days of handling for ∼5 min each day, followed by 1–2 days of habituation to the optical tether in their home cage for 10 min. All behavior video was captured via a USB webcam (Logitech) connected to a computer running ANY-maze (Stoelting) which was used to track the position of the mouse during behavior. Mice were randomly assigned to a viral condition and the experimenter was blinded to the mouse’s virus assignment during all subsequent behavioral assessment or analyses.

### Tail suspension test (TST)

TST mice were suspended by taping their tail to a bar ∼12 inches above the tabletop, within 4 inch wide stalls of a custom apparatus constructed for this purpose. A 1.5 ml microcentrifuge tube was cut and placed over the tail base to prevent the mouse from climbing its tail during testing. The TST for optogenetics experiments was 14 minutes in duration and only one mouse was run at a time. Optogenetic stimulation was delivered in 4x, 3 minute blocks (OFF-ON-OFF-ON) with stimulation parameters as per below. For all other experiments, the TST was 12 minutes in duration and up to 4 mice were run simultaneously in separate stalls where they were not visible to each other. The apparatus was thoroughly cleaned between uses. The first 2 minutes of the TST were discarded as per standard protocols and analyses were conducted using videos of behavior during the TST (see, (21)). TST videos for VTA terminal optogenetics were analyzed manually on a per minute basis by an experimenter that was blinded to the conditions being tested and using an established method (21) For TST videos from Drd2 deleted mice, we developed an unbiased, automated analysis algorithm using Matlab, that allowed us to quantify not only total duration of struggling, but also the number and duration of individual struggling episodes. First, we developed a custom analysis process where previously collected videos were analyzed using ANY-maze software (Stoelting). A region of interest was drawn around the base of the mouse’s tail in order to track lateral movement during the TST. Several points of tracking were assessed as potential measures of struggling/immobility, but tailbase tracking most faithfully captured distinct episodes (data not shown). The output of this process was a sequence of x-y coordinates corresponding to tailbase position at each frame of the recorded behavior video. These coordinates were processed via a custom Matlab algorithm to produce a smoothed, average velocity in the lateral direction for every frame. We then used a supervised machine learning algorithm that used human annotated videos to train an algorithm to classify struggling and immobility epochs from tailbase velocity values. Additional Matlab code was used to calculate the number and duration of individual episodes and compare between conditions.

### Open field (OF)

OF testing was conducted in a standard sized arena (40×40cm), with a height of 30cm. Video was collected from above and subsequent analysis was done using built-in ANY-maze tracking. Open field testing was 8 minutes in duration and mouse was placed by hand into the lower right corner of the apparatus. Optogenetic stimulation was delivered in 4x, 2 minute blocks (OFF-ON-OFF-ON) with stimulation parameters as per below. Total distance traveled in the apparatus, total or on a per minute basis, was analyzed as a measure of overall activity level.

### Social Defeat Stress

Social defeat stress experiments were conducted consistent with standardized protocols for this procedure (22). Retired, singly housed CD-1 male breeder mice (Charles River; 4-6 months of age) were screened for adequate aggression prior to starting protocol. Male C57 BL/6 mice between 5-6 weeks of age were exposed to 10 minutes of defeat from CD-1 aggressor mice, followed by 6-8 hours of exposure to the aggressor in a cage divided by a perforated plexiglass barrier for 10 consecutive days. Aggressor mice were rotated each day to prevent habituation and the quantity and quality of aggressive episodes was monitored to ensure even exposure to defeat. Control mice were exposed similarly, but to cagemates instead of aggressor mice. 24 hours after the last defeat, mice were tested for social interaction in a standard open field arena in which a completely novel CD-1 mouse was secured inside a wire-mesh cup. Mice were briefly exposed to the CD-1 mouse, followed by exposure to an empty cup alone (3 minute each, separated by 30s). A social interaction zone around the cup was established and social interaction ratio (SI ratio) calculated as time spent in the interaction zone with the CD-1 mouse divided by without. SI ratios for defeated mice roughly segregated into two clusters, those within the range of control mice were defined as ‘resilient’ and those much lower as ‘susceptible’ for the purposes of our experiment. We then cut slices from resilient and susceptible littermate mice within 48 hours of the social interaction test as per protocol described above. Susceptible and defeated mice were imaged on the same in counterbalanced order to ensure consistent time between the last defeat the imaging experiment.

### Optogenetic stimulation protocols

For optogenetic stimulation during behavior, light stimulation was delivered via fiber optic cable fed through a commutator (Doric Lenses) and attached to a 100 mW, 473 nm laser (OEM). The laser was driven with a pulse generator using two protocols: In the tonic protocol a 5 Hz train of 4 ms pulses was used while in the phasic protocol 500 ms bursts of 50 Hz 4 ms pulses were delivered every 5 seconds. The total light power delivered during a pulse was 5 mW.

### Statistics

Unless otherwise specified, nonparametric tests or ANOVA was used to assess significance. Statistics were calculated using custom MATLAB code or Graphpad Prism. All statistical parameters, including test statistics, correction for multiple comparisons, and sampling of repeated measurements are stated in the main text. All tests are two-sided, and error bars represent SEM. The code and data used to generate and support the findings of this study are available from the corresponding author upon reasonable request.

## RESULTS

### Measuring network states in a slice GCaMP assay

To examine putative neural signatures associated with antidepressant treatment and/or depression susceptibility in prefrontal circuits, we targeted viral expression of GCaMP6 to deep layer prefrontal neurons and observed activity patterns generated in isolated microcircuits from live prefrontal slices (**Figure 1A-D**). A standard approach to analyzing GCaMP imaging experiments has been to infer neuronal firing patterns based on large ‘events’, which are presumed to reflect bursts of action potentials. However, a more conservative approach is to avoid assumptions about the relationship between GCaMP fluorescence and underlying electrical activity, and directly analyze the fluorescence signals. One reason that most studies focus on events, rather than GCaMP fluorescence signals themselves, is that GCaMP signals obtained via single-photon imaging may be contaminated by shared background fluorescence. This is potentially problematic when examining patterns of circuit activity, rather than single cell activity, because correlations between signals from different regions of interest (ROIs), intended to measure activity in different cells, may end up being dominated by autocorrelation between shared background fluorescence. To overcome this issue, we computed correlations between fluorescence in one cell and the *derivative* of fluorescence from other cells. Because the integral of a signal with its derivative is zero over a closed path, this will minimize autocorrelation driven by shared background fluorescence – indeed we have previously verified that this is the case and used this technique to examine anxiety-related activity in vivo (20). Intuitively, correlation computed this way measures the degree to which a GCaMP signal in ROI *i* predicts increases or decreases in ROI *j*. The matrix of correlations between all sources represents an abstraction of the network state during a given window of time (**Figure 1D**). This relatively simple method maximizes the use of rich GCaMP signals to define a metric for the organization of activity in a microcircuit.

### Antidepressants and D2Rs enhance correlated circuit activity

First, we studied the effects of two clinically effective antidepressant medications, which have distinct mechanisms of action and effect profiles: the selective serotonin reuptake inhibitor (SSRI) fluoxetine, which is only effective after chronic treatment (e.g., for weeks), and the N-methyl-D-aspartate Receptor (NMDAR) antagonist, ketamine, which rapidly alleviates depression (within hours), including in patients with treatment-refractory or bipolar depression (23–27). Prefrontal slices from mice that received either chronic fluoxetine (daily injections for 3 weeks) or a single dose of ketamine both exhibited a marked increase in positively correlated activity between mPFC neurons (**Figure 1E,F**). Next we measured correlations after activating dopamine D2 receptors (D2Rs). Repeated stress can decrease the firing of dopaminergic neurons which project to prefrontal cortex (28), as well as dopamine release in the prefrontal cortex (29). This has led to the hypothesis that deficient prefrontal dopamine may contribute to depression (29–31). Indeed, D2R agonists such as pramipexole are effective antidepressants that modulate prefrontal circuits (32,33), and D2Rs have been implicated in the therapeutic effects of both selective serotonin reuptake inhibitors (SSRIs) and exercise (34–38). Therefore, we hypothesized that prefrontal D2R activation might elicit antidepressant-like circuit effects, and conversely, that manipulations related to depression might alter network responses to D2R activation. Indeed, application of the D2R agonist quinpirole (10 µM) elicited an increase in positive correlations (**Figure 1G**), similar to what we had observed in fluoxetine or ketamine-treated mice.

### Models of depression susceptibility have deficits in D2R-driven correlated activity

We also wondered whether mice which model aspects of depression might exhibit the opposite changes – reduced positive correlations either at baseline, or in response to D2R activation. Chronic social or psychological stress is a major risk factor for the development of depression in patients (39). In mice, this can be modeled in an ethologically relevant way by chronic social defeat stress, in which mice are repeatedly exposed to social dominance interactions from a larger, more aggressive mouse (22). Mice exposed to chronic social defeat exhibit a continuum of responses. These mice can be classified as either “susceptible” or “resilient” based on their behavior in an assay measuring social avoidance of a novel mouse. All mice tend to develop stress and anxiety-like responses to social defeat, but only susceptible mice, i.e., mice which avoid the novel mouse, exhibit more extensive, depression-like changes in behavior (22,40). We obtained prefrontal slices from mice which had undergone social defeat and been classified as either susceptible or resilient, then imaged prefrontal circuit activity before and after applying quinpirole to activate D2Rs (**Figure 2A-E**). Slices from resilient and susceptible mice had similar positive correlations at baseline, but exhibited strikingly different responses after D2R stimulation (**Figure 2F,G**). After quinpirole application, positive correlations in resilient mice were increased several-fold compared to susceptible mice.

**Fig. 2.**
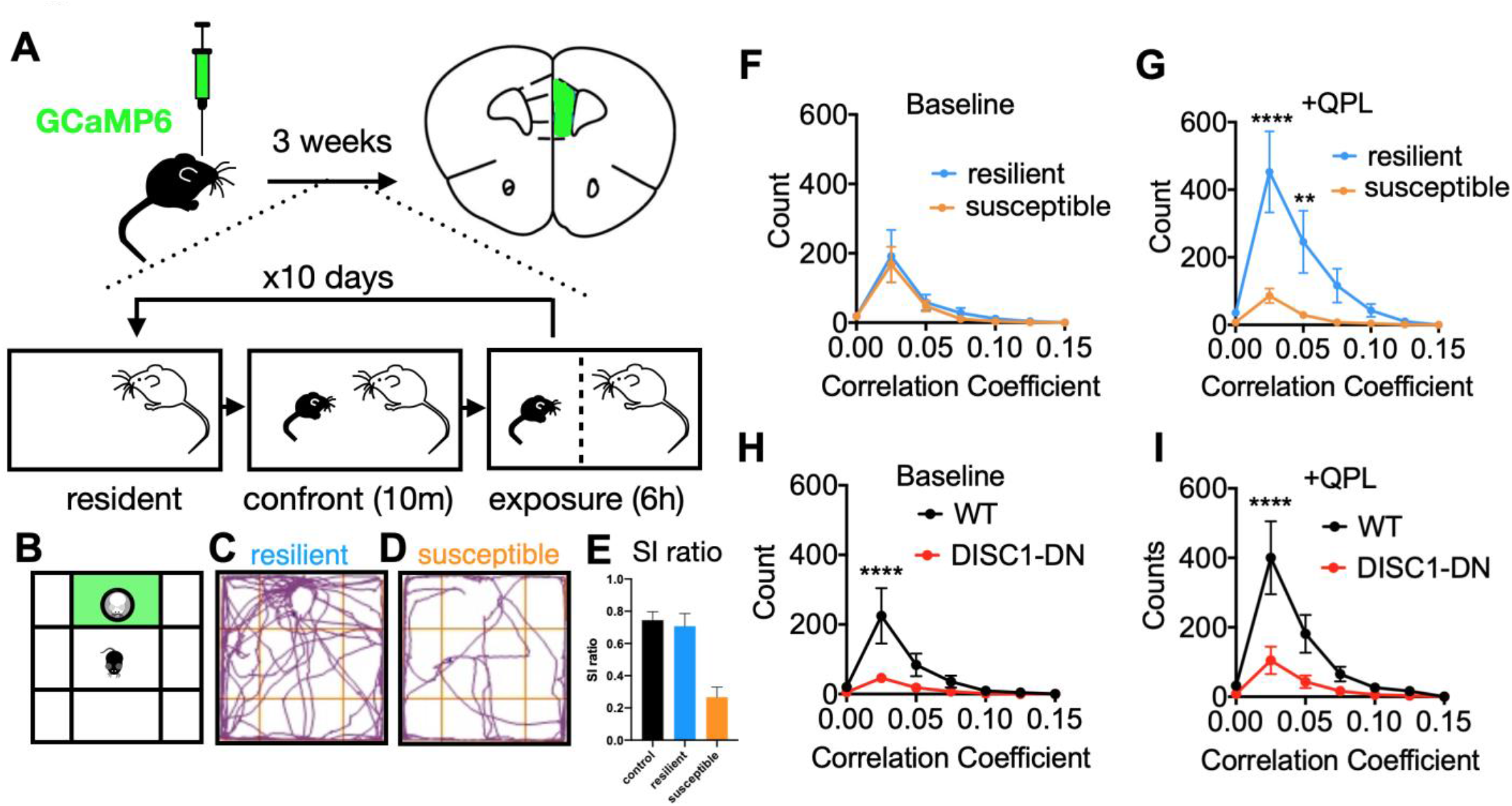
The effect of quinpirole on correlated activity is blunted in mouse models of depression susceptibility. (**A**) Schematic of injection and social defeat stress protocol. (**B**) Social interaction assay with green interaction zone, (**C**) social behavior resilient mouse, (**D**) social behavior susceptible mouse. (**E**) social interaction ratio scoring (time in interaction zone with/without novel CD1). (**F-G**) Comparison of positive correlations in slices prepared from resilient and susceptible mice during (**F**) baseline conditions [*F*_Interaction_(6,63)=0.064, *P*=0.999, *F*_Group_(1,63)=0.421, *P*=0.519] and (**G**) after Quinpirole wash-in (20-40min) [*F*_Interaction_(6,63)=5.99, *P*<0.0001, *F*_Group_(1,63)=27.2, *P*<0.0001]. (**H-I**) Comparison of positive correlations in slices prepared from wildtype (WT) and DISC1-DN mice during (**H**) baseline conditions [*F*_Interaction_(6,98)=2.77, *P*=0.016, *F*_Group_(1,98)=8.64, *P*=0.004] and (**I**) after Quinpirole wash-in (20-40min) [*F*_Interaction_(6,98)=3.81, *P*=0.002, *F*_Group_(1,98)=14.31, *P*=0.0003], 2-way ANOVA with Sidak post-hoc test, N=6,5,9,7 for susceptible, resilient, WT, DISC1-DN, respectively.

Environmental influences such as stress are often causal or contributory in clinical depression, however heritability accounts for approximately 37% of risk for developing major depressive disorder (10,39). A rare, but highly penetrant truncation in the gene *Disc1* was originally associated with major mental illness, including schizophrenia, bipolar disorder, and depression, in a Scottish pedigree (41–43). Genetic studies have found evidence for an association between specific *Disc1* variants and recurrent major depression, though these studies notably lack the sample sizes needed for genome-wide significance (44). Nevertheless, many mouse models with disruptions in *Disc1* exhibit specific changes in behavior similar to those seen in other models of depression and opposite to those elicited by antidepressant treatments (11,45–47). Therefore, we studied one of these models to determine whether genetic factors which influence responses to stress might also exhibit altered correlated activity in prefrontal circuits. We found that mice expressing a dominant negative *Disc1* variant (*Disc1-DN*), had significantly reduced positive correlations in our slice assay, both before and after D2R activation (**Figure 2H,I**). This deficit in positively correlated activity may reflect changes in circuits that produce alterations in behavior similar to those seen in other depression models and opposite to those produced by antidepressant treatment, e.g., increased immobility in the TST.

Depression is highly heterogeneous, such that individual assays and models typically capture only a portion of this complex syndrome. That being said, certain underlying mechanisms may represent points of convergence that are relevant to diverse behaviors and manipulations, even if they do not capture the entirety of the disorder. The preceding results suggest a simple model in which intrinsically generated, positively correlated activity between prefrontal neurons represent one such point of convergence. In this simple model, antidepressant manipulations and stress-resilience either directly increase intrinsically generated correlated activity between prefrontal neurons, or augment this activity in response to D2R activation. By contrast, in stress-susceptible mice and mice that exhibit increases in inactive responses to stress, this correlated activity is decreased.

How can we assay whether changes in correlated activity between prefrontal neurons are relevant for behaviors that are impacted by stress and/or reflect aspects of depression? Although the role of PFC in depression is incompletely understood, cognitive dysfunction is a core feature and may predict poorer outcomes (48). Amongst these, deficits in adaptively responding to challenging circumstances, flexible decision making, and cognitive bias are common (48). A commonly used assay to measure the coping behavior of rodents in response to acute stress is the tail suspension test (TST). Tail suspension confronts a mouse with an inescapable stressor and mice respond by struggling to escape (active coping) or adopting an energy conserving immobile posture (passive coping). Importantly, the TST has largely failed as a screen for discovering new antidepressants, because it is highly sensitive to compounds which recapitulate the mechanisms of action of existing antidepressants. Furthermore, aside from face validity, there is no bona fide reason to think of immobility as an analog of the depressed state. That being said, a striking observation is that all known antidepressant treatments including both SSRIs and ketamine increase struggling in the TST. Moreover, both *Disc1* mutant mice and mice exposed to social defeat stress have been shown to exhibit decreased struggling in the TST (49). This suggests that the disparate molecular and cellular pathways engaged by antidepressants (SSRIs and ketamine) and models of depression (*Disc1*-DN and social defeat) ultimately converge on common neural mechanisms that result in changes in neural activity that manifest within the TST, even if the original rationale for this assay (face validity) is no longer accepted.

### Phasic dopamine release in mPFC promotes effortful responses to acute stress

While the relationship between the TST (or related assays) and depression has been challenged, its construct validity for coping with adversity is stronger (50–52). Mesolimbic dopamine as well as the manipulations investigated in our ex vivo assay affect behavior in the TST, but the role of prefrontal dopamine is less clear (28,53). Generally, D1 and D2Rs are expressed by distinct neuronal populations and often regulate opposing cellular, circuit and behavioral functions (54–58). VTA dopamine neurons fire either in a low frequency, sustained manner or in high frequency phasic bursts which preferentially release dopamine (59–62). We have shown that in PFC, phasic activation of VTA inputs release high concentrations of DA (12), which may preferentially elicit a D2R-driven network state associated with labile neural activity that underlies cognitive and behavioral flexibility (12,57,63). Because D2R stimulation elicits a positively correlated state that is also associated with stress-resilience and antidepressant response (**Figures 1** and 2), we hypothesized that phasic stimulation would enhance struggling in the TST. To test this hypothesis, we targeted channelrhodopsin-2 expression to VTA dopaminergic neurons and activated terminals in mPFC by delivering blue laser light using a fiber optic implant (**Figure 3A–C**). To model distinct tonic vs. phasic firing patterns observed *in vivo*, we used either a continuous 5 Hz pattern of light pulses (tonic) or arranged the same number of pulses into 50 Hz bursts occurring every 5 seconds (**Figure 3D,** see also (12)). Phasic, but not tonic, stimulation of prefrontal DA release increased effortful responses (i.e., struggling) specifically during light-on periods (**Figure 3E,F**). Neither phasic nor tonic stimulation produced nonspecific increases in locomotion for mice exploring an open field (**Figure 3G,H**). This indicates that phasic release specifically enhances effortful responses to stress, not generic motor activity.

**Fig. 3.**
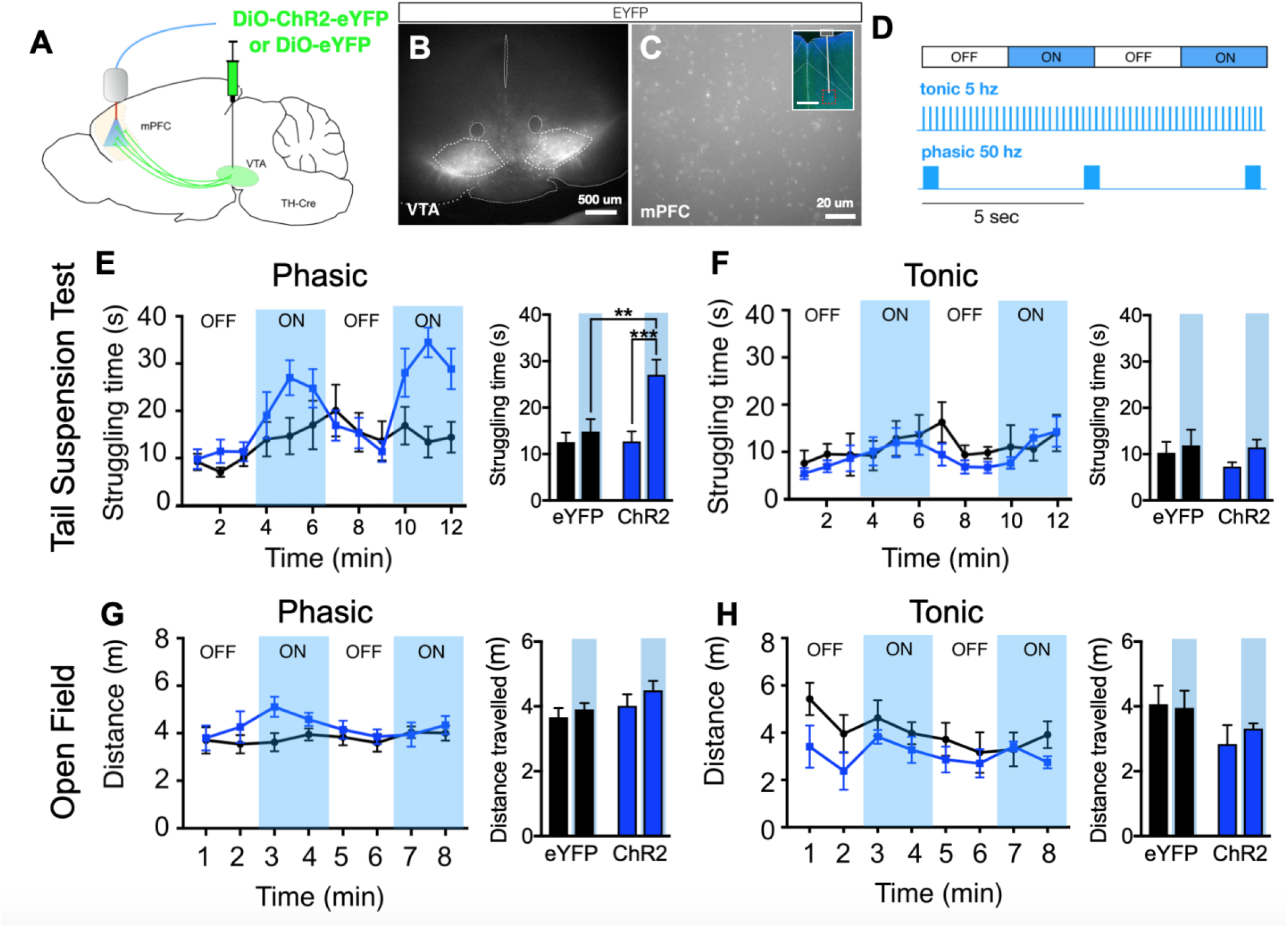
Phasic dopamine stimulation in mPFC increases struggling time during the tail-suspension test. (**A**) Experimental design. (**B**) Representative image of ChR2-eYFP expression in the ventral tegmental area (VTA). (**C**) eYFP+ terminals in mPFC (inset shows fiber tract). (**D**) Optogenetic stimulation protocols. (**E–F**) Quantification of struggling time during the tail suspension test during (**E**) phasic stimulation of VTA terminals [*F*_*Interaction*_(1,36)=5.23, *P*=0.28, *F*_*Laser*_(1,36)=9.74, *P*=0.004, *F*_*Virus*_(1,36)=5.43, *P*=0.023], or (**F**) tonic stimulation of VTA terminals [*F*_*Interaction*_(1,24)=0.391, *P*=0.538, *F*_*Laser*_(1,24)=1.893, *P*=0.182, *F*_*Virus*_(1,24)=0.686, *P*=0.416]. (**G– H**) Quantification of distance travelled in an open field during (**E**) phasic stimulation of VTA terminals [*F*_*Interaction*_(1,34)=0.179, *P*=0.683, *F*_*Laser*_(1,34)=1.511, *P*=0.228, *F*_*Virus*_(1,34)=2.604, *P*=0.116], or (**F**) tonic stimulation of VTA terminals [*F*_*Interaction*_(1,22)=0.365, *P*=0.552, *F*_*Laser*_(1,22)=0.139, *P*=0.713, *F*_*Virus*_(1,22)=3.646, *P*=0.069]. Blue shading indicates laser on periods. 2-way ANOVA with posthoc Sidak test, N=9,11,10, eYFP, ChR2 (TST), ChR2 (OF) respectively.

### Prefrontal D2R deletion shortens effortful responses to acute stress

If, as hypothesized, the phasic release of prefrontal dopamine increases struggling by acting on D2Rs, then deleting D2Rs from mPFC may have an opposing effect on behavior in the TST. To test this hypothesis, we injected the bilateral mPFC of floxed D2R (Drd2^loxP/loxP^) mice with adeno-associated virus expressing Cre recombinase or a control virus and after allowing time expression/deletion conducted the TST and OF tests (**Figure 4A-C**). In order to analyze the fine structure of effortful responses to tail suspension stress, we developed an unbiased, automated algorithm that uses tailbase velocity to assess both the number and duration of individual struggling epochs (**Figure 4D-F**). Prefrontal-specific deletion of D2Rs had no significant impact on the overall time spent struggling, but mice with D2Rs specifically deleted from mPFC exhibited a greater number of struggling episodes which were shorter in duration (**Figure 4G-I**). Deletion of mPFC D2Rs did not result in any change in overall activity level as measured by distance traveled in the OF assay (**Figure 4J,K**). These results are consistent with the hypothesis that prefrontal dopamine release, elicited by phasic firing patterns, act via D2Rs to drive a positively correlated activity state that is downstream of both antidepressant treatments and factors promoting depression-like states or depression susceptibility. This dopamine-influenced prefrontal state is a plausible neural mechanism that could support the allocation of effort or maintenance of active stress-coping more generally (see model, **Figure 5**). In summary, we find that depression-related manipulations may alter the organization of prefrontal activity to bias behavioral responses during challenging situations. Such a mechanism is consistent with the cognitive model, where negative perceptual and cognitive bias are central to development and maintenance of depressed states (48).

**Fig. 4.**
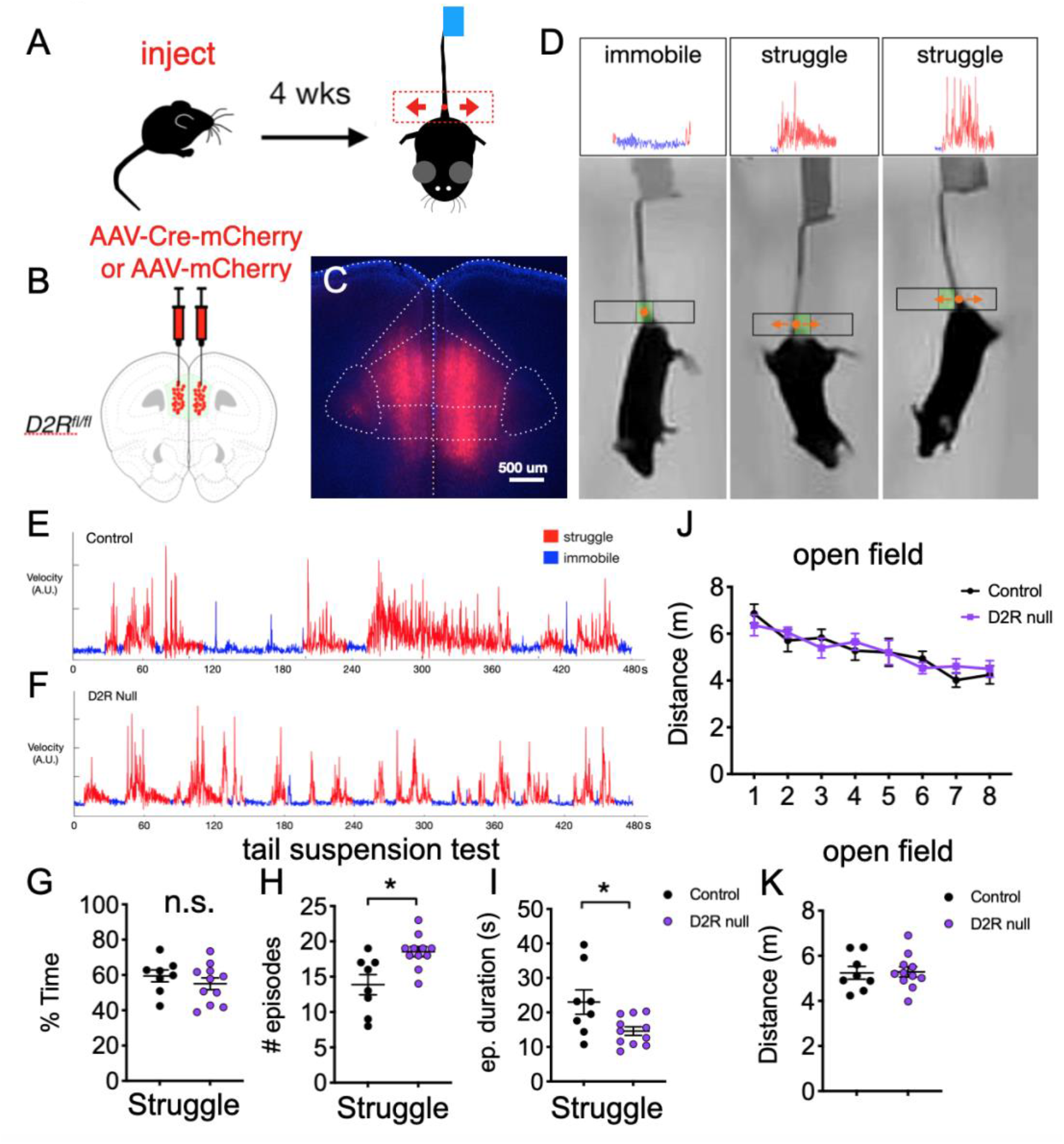
Dopamine D2 receptor deletion reduces duration of struggling episodes during TST. (**A**) Experimental design. (**B**) Schematic for injection of Cre expressing vs. control virus into mPFC in D2R flox/flox mice. (C) Representative image showing bilateral Cre-mCherry expression limited to mPFC. (D) Representative video still shots of immobile vs. struggling mice engaged in TST with tailbase velocity tracking of episode shown above. (E-F) Representative examples of tailbase velocity tracking for control vs. D2R deleted mice with struggling and immobile epoch classifications shown. (G-I) Quantification of (G) total struggling time, (H) number of distinct struggling episodes and (I) mean duration of individual struggling episodes. (J-K) Quantification of distance traveled in open field test. Unpaired t test, n.s. = not significant; * p<0.05.

**Fig. 5.**
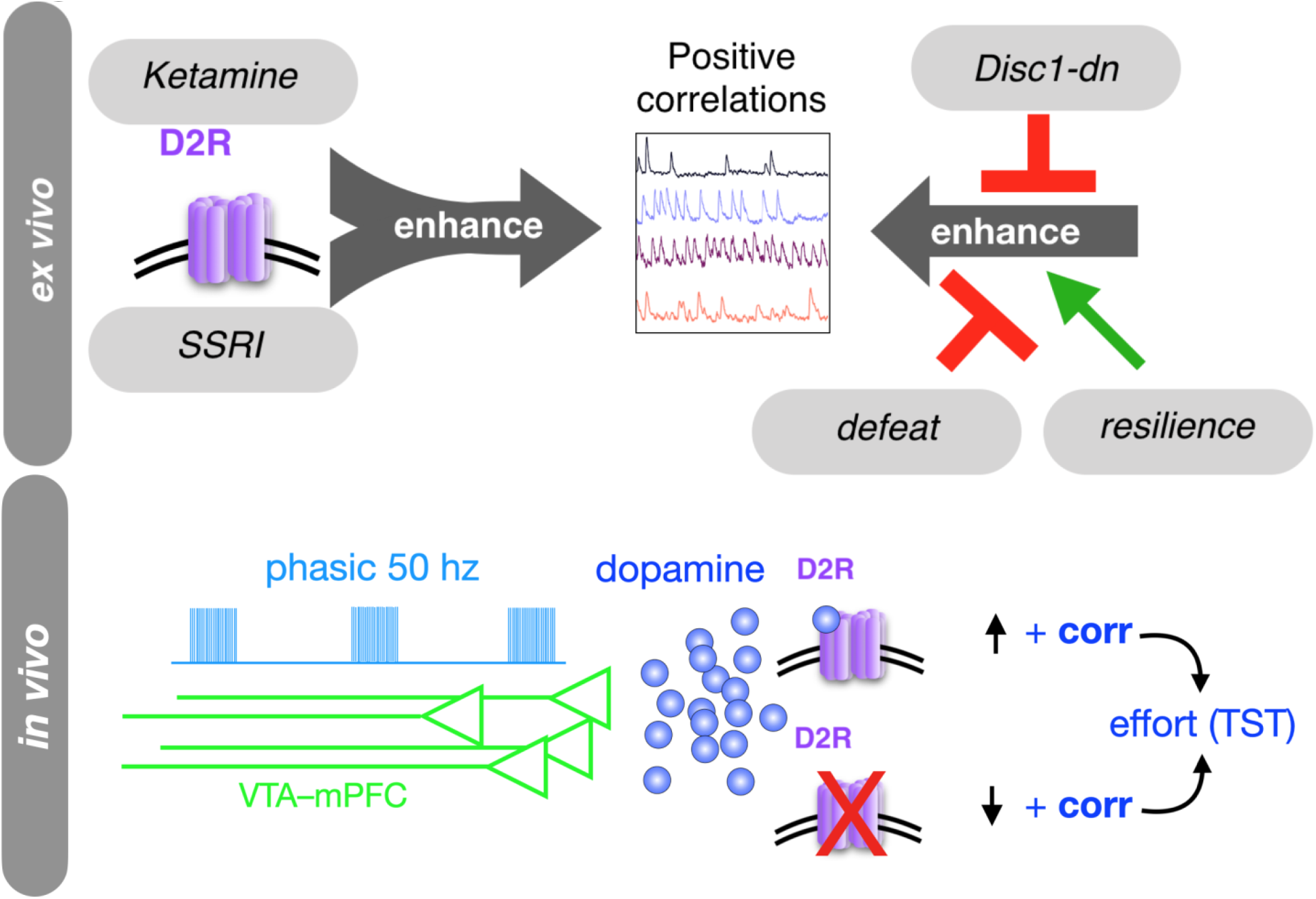
Model showing the convergent effects of D2Rs, antidepressants, and depression susceptibility models on positively correlated mPFC activity, *ex vivo*. Model also proposes that dopamine – via phasic bursting of VTA-mPFC afferents – acts via D2Rs to induce changes in effortful coping by recruiting similar prefrontal network states.

## DISCUSSION

Previous studies have largely focused on using single preclinical models to study neural and behavioral correlates of purported importance for depression. However, a given manipulation can have pleiotropic effects, many of which may not be relevant for the clinical condition being studies. This is particularly pertinent because depression is heterogeneous and existing models do not perfectly recapitulate the human condition. However, the bidirectional convergence of multiple manipulations associated with both antidepressant treatment and depressional susceptibility onto a common neural process makes it more likely that this process is relevant to the shared clinical entity. Here, we demonstrate a new approach, using calcium imaging to reveal convergence of genetic, environmental and pharmacological manipulations related to depression onto patterns of correlated activity in prefrontal microcircuits (**Figures 1** and 2). Then we used optogenetic stimulation and targeted gene deletion to confirm that prefrontal dopamine signaling modulates not only these patterns of activity, but also behaviors that are commonly used to assay antidepressant efficacy and depression susceptibility (**Figures 3** and 4). This work describes a general, discovery-oriented framework for identifying neural signatures relevant to disease and validating their relationships with behavior *in vivo*.

Multiple lines of evidence implicate prefrontal dopamine in the response to acute and chronic stress, as well as aversive processing more generally (64,65,67). D2Rs are less well studied, but may underlie resolution of depression, for example secondary to exercise or antidepressant treatment (34–38). Our study takes this several steps further by linking D2Rs to a specific change in patterns of correlated activity amongst prefrontal neurons. Rhythmic neuronal activity is important for local computations and couple prefrontal activity with activity in connected regions important for behavioral deficits in depression (66). For example, theta and low gamma synchrony in mPFC is disrupted in Disc1 mutant mice and correlates with active coping behavior in the TST (49). While the relationship between the correlated patterns of activity we studied and other neural processes including rhythmic synchronized are currently unknown, one intriguing possibility is that positive correlations reflect the tendency for mPFC microcircuits to enter particular modes of synchronized rhythmic firing which are engaged during stress coping. Patterns of correlated prefrontal activity might also depend on the formation of postsynaptic dendritic spines on prefrontal projection neurons, a process that has recently been shown to be enhanced by ketamine, suppressed by stress-related manipulations, and involved in sustaining antidepressant-driven increases in struggling in the TST (68).

Psychological stress worsens most psychiatric conditions and its effects on the PFC in particular may disrupt cognitive processes such as decision making (69). Prefrontal dopamine signaling, particularly phasic dopamine release and D2R activation have been implicated in cognitive and behavioral flexibility (12,70–72). Flexible decision making or more generally, adaptation to changing task demands under stress could tie together our findings as part of an intermediate phenotype. In this model, positive correlations could be a signature of general cognitive flexibility. Phasic DA release and D2R activation may drive a prefrontal state necessary for a mouse to adopt or maintain active coping strategies, rather than becoming excessively biased towards an immobile state. VTA neurons are thought to encode reward prediction errors via phasic bursts occurring after unexpected positive outcomes or rewards (73). Thus, such a signal indicates that a situation is better than expected in some way and can trigger associated learning or open the system to new action-outcome associations (74,75). Imposing a phasic signal during passive coping may trigger struggling by altering the real-time assessment of possible outcomes, e.g., the probability of escape. Given that VTA neurons signal something about positive versus negative outcome predictions, an intriguing possibility is that positively correlated activity is more broadly related to the extent to which prefrontal circuit are biased towards predicting positive versus negative outcomes. Depression is characterized by cognition that is biased towards negative outcomes, so antidepressants may alter this balance via their effect on correlation state. The particular biomarker identified here will require further characterization and validation, but is an intriguing candidate mechanism for linking dopaminergic signaling and prefrontal activity with stress-induced pathological states and their resolution.

## Acknowledgements

We wish to thank Dr. Laura DeNardo, Dr. Nicholas Frost and members of the Sohal lab for helpful comments and discussion related to the manuscript.

## Data and materials availability

All code and data used in the analysis is available to academic researchers upon reasonable request.

